# Fingerprinting and behavioural prediction rest on distinct functional systems of the human connectome

**DOI:** 10.1101/2021.02.07.429922

**Authors:** Maron Mantwill, Martin Gell, Stephan Krohn, Carsten Finke

## Abstract

The prediction of inter-individual behavioural differences from neuroimaging data is a rapidly evolving field of research, focusing on individualised methods to describe human brain organisation on the single-subject level. One method that harnesses such individual signatures is functional connectome fingerprinting, which can reliably identify individuals from large study populations. While connectome fingerprints have been previously associated with individual cognitive function, these associations rest on indirect evidence.

Contrasting with these previous reports, here we systematically investigate the link between connectome fingerprints and the prediction of behaviour on different levels of brain network organisation (individual edges, network interactions, topographical organisation, and edge variability), using 339 resting-state fMRI datasets from the Human Connectome Project.

Our analysis revealed a significant divergence between connectivity signatures that discriminate between individuals and those predictive of behaviour on all levels of network organisation. Across different parcellation schemes, thresholds and prediction algorithms, we consistently find fingerprints in higher-order multimodal association cortices, while neural correlates of behaviour display a more variable topological distribution. Furthermore, we find the standard deviation of connections between subjects to be significantly higher in fingerprinting than in prediction, making inter-individual connection variability a possible separating marker.

These results demonstrate that participant identification and behavioural prediction involve highly distinct functional systems of the human connectome, suggesting that connectome fingerprints are not as functionally relevant as previously believed. The present study thus calls for a re-evaluation of the significance of functional connectivity fingerprints in personalized medicine.

## Introduction

The mapping of individual cognitive and behavioural performance to neurological patterns, and the identification of robust disease biomarkers are primary goals of neuroscience (Castellanos, Di Martino, Craddock, Mehta, & Milham, 2013; Woo, Chang, Lindquist, & Wager, 2017). Indeed, the ability to predict individual cognitive performance or a subject’s disease progression is regarded as a prerequisite for the development of a personalized medicine (Gabrieli, Ghosh, & Whitfield-Gabrieli, 2015; Eickhoff & Langner, 2019). This focus on the individual requires a shift from group-level to single-subject analyses, moving the focus from finding average differences between groups into a more mechanistic understanding of the underlying processes (Finn et al., 2017).

Connectome fingerprinting represents one such individualised and powerful approach to single-participant analysis has been viewed as a way to map individual differences in functional organisation onto individual cognition and behaviour (Finn et al., 2015, 2017; Emerson et al., 2017; Waller et al., 2017; Liu, Liao, Xia, & He, 2018; Horien, Shen, Scheinost, & Constable, 2019). In connectome fingerprinting individual participants can be reliably identified within large datasets with accuracies exceeding 90%, based on the discriminatory power of individual functional connectomes. Interestingly, it has been reported that the resting-state networks that best discriminate between individuals are the same networks that are predictive of individual differences in cognitive performance and behaviour (Finn et al., 2015). However, the assumption that networks that best discriminate between individuals are also behaviourally relevant is only based on visual inspection of the similarity of networks central for identification and prediction. Thus, a robust statistical analysis of the link between the connectivity patterns contributing to subject discrimination and behavioural prediction is still missing.

Here, we investigate the relevance of connectome fingerprinting to behavioural prediction. We replicate the analysis from Finn et al. (2015), yielding the suggestive overlap based on visual inspection. However, a systematic examination of the overlap of the respective network patterns shows that discriminatory connectivity signatures and connections predictive of behaviour are unrelated, both on single-edge and network levels as well as differ in their topographical distribution. These findings are robust with respect to different parcellation schemes, psychometric variables and prediction algorithms. Together, our results suggest an alternative perspective on the relation between fingerprinting and behavioural prediction that rests on edge-level variability.

## Methods

### Dataset

We used the unrelated subjects sample (n = 339, 156/183 m/f, ages 22-35) from the full release of the publicly available Human Connectome Project dataset (Van Essen et al., 2013). In our prediction analysis, we excluded subjects that had missing behavioural data in a case-by-case fashion (Table 1). The HCP scanning protocol was approved by the local Institutional Review Board of Washington University in St. Louis, MO, USA, the details of which are described elsewhere (Van Essen et al., 2013). Briefly, for resting state fMRI (rs-fMRI), whole-brain multiband gradient-echo-planar images were acquired on a 32-channel 3T Siemens “Connectome Skyra” scanner with TR = 720 ms, TE = 33.1 ms, flip angle = 52 degrees, bandwidth = 2,290 Hz/pixel, in-plane field of view = 208×180 mm^2^, 72 slices, 2 mm isotropic voxels and 1,200 volumes (14 min and 24 s). Rs-fMRI sessions were acquired left-to-right (LR) and right-to-left (RL).

**Table 1.** Results for Psychometric Prediction

| Psychometric Variable | Spearman Correlation | *p*-values | Subjects (*n*) |
| --- | --- | --- | --- |
| Fluid cognition composite score | 0.22 | .002 | 318 |
| Crystallised cognition composite score | 0.21 | .001 | 320 |
| Total cognition composite score | 0.25 | < .001 | 318 |
| Cognitive flexibility | 0.18 | .006 | 319 |
| Fluid Intelligence | 0.22 | < .001 | 319 |
| Sustained Attention (Specificity) | 0.17 | .015 | 319 |
| Grip strength | 0.44 | < .001 | 319 |
| Dexterity | 0.23 | < .001 | 320 |
| Language/Reading Decoding | 0.19 | .003 | 320 |
| Language comprehension | 0.19 | .004 | 320 |
| Spatial Orientation | 0.21 | .002 | 319 |
| Emotion Recognition | 0.20 | .002 | 319 |

Last row contains the number of subjects with complete data. All *p*-values are permutation-derived and FDR corrected for all behavioural predictions.

### rsfMRI preprocessing

We closely followed Finn et. al. (2015) in our pre-processing pipeline and used the minimally preprocessed rs-fMRI dataset (Glasser et al., 2013). This included gradient distortion correction, motion correction, image distortion correction, registration to MNI standard space and intensity normalization. We then used the CONN toolbox (Whitfield-Gabrieli & Nieto-Castanon, 2012) for SPM12, regressing out 12 motion parameters (provided with the HCP dataset under Movement_Regressors_dt.txt), mean time courses of white matter, CSF and the global grey matter signal (approximating global signal). Linear trend was removed and the data were band-pass filtered (0.01 – 0.1Hz). We did not perform any smoothing. The resulting voxel-wise time series were parcellated using four different atlases: for the main analyses, we used the Shen atlas with 268 nodes (i.e., regions of interests (ROIs)) that was also used in Finn et al. (2015); for validation, we used the Brainnetome atlas (Fan et al., 2016), the HCP MMP 1.0 atlas (Glasser et al., 2016), and the AAL atlas (Tzourio-Mazoyer et al., 2002). For every parcellation scheme, we extracted the nodal time series by averaging over all voxels within the respective ROI.

The Shen atlas is a whole-brain atlas derived from functional connectivity and defined using a group-wise spectral clustering algorithm. The three other atlases cover different levels of detail as well as different approaches to the definition of nodes. The AAL atlas provides an anatomy-based parcellation with 90 cortical and 26 cerebellar nodes. The Brainnetome Atlas (Fan et al., 2016) is a whole-brain atlas containing a similar number of nodes to the Shen atlas (210 cortical and 36 subcortical nodes) and is defined using both anatomical and functional connections. HCP MMP 1.0 is a very detailed cortical in-vivo parcellation with 360 nodes. We acquired resting-state network definitions for all four atlases. The Shen and Brainnetome atlases provide resting-state network definitions for each node, and for the later these are based on the established Yeo-7 resting-state networks (Yeo et al., 2011). Since the Yeo-7 network definition does not assign subcortical nodes to a network, we created an eighth subcortical network. For the HCP MMP 1.0, we relied on network assignments supplied by (Ji et al., 2019), partitioning the 360 nodes into 12 resting-state networks. The AAL nodes were split into 5 resting-state networks based on network definitions from (He et al., 2009). Here, we created a sixth cerebellar resting-state network including all cerebellar nodes.

### Functional connectome

Individual functional connectomes were built as functional connectivity matrices calculated as the Pearson correlation between the time courses of all region-to-region pairs. In the framework of functional brain networks (Rubinov & Sporns, 2010), each ROI represents a network node, and the connection between two ROIs represents an edge in the network.

Each participant has two resting state scans (LR and RL encoding) by session, thus creating four functional connectivity matrices by individual. We averaged the two matrices from one session (LR and RL), resulting in two final matrices (one per session) for every individual (matrix dimension: number of nodes by number of nodes, the exact number depending on the parcellation scheme). The final functional connectivity matrices were z-scored, and the upper triangle was vectorized.

### Functional connectome fingerprinting

Fingerprinting was performed as in Finn et al. (2015). Briefly, the functional connectome of a ‘source’ subject at timepoint t1 is used to identify the same subject at timepoint t2, referred to as ‘target’. The target session is identified from a pool of functional connectomes containing both the target connectome as well as connectomes of other ‘distractor’ subjects. Identification is performed by correlating the FC vector of the ‘source’ from one of the two scanning sessions (e.g., “rest 1”) with the FC vectors of all 339 subjects (including the ‘target’) in the other session (e.g., “rest 2”), resulting in 339 correlations. The subject with the highest correlation coefficient is picked and assigned a score of 1 if the picked subject matches the target identity (hit), and a score of 0 otherwise (miss). This procedure was applied for all possible session 1 to session 2 source-target pairs (i.e., 339 identifications) and then repeated once more for all session 2 to session 1 source-target pairs (again 339 identifications). Lastly, we performed a nonparametric permutation test with 1000 permutations to examine the statistical significance of our identification analysis. In each permutation, the target participant’s and distractor session’s identity were randomized and fingerprinting accuracy recorded. P-values were then calculated as the proportion of randomly permuted instances exceeding the empirically observed accuracy over all permutations.

### Network and edge contributions to fingerprinting

To assess the contribution of different resting-state networks, we calculated the differential power (DP) of different edges (i.e., node-to-node connections), using publicly available scripts (Finn et al., 2015). DP of an edge reflects an edge’s “uniqueness” and stability and thus its ability to differentiate an individual. First, we exclusively investigated the DP of within-network edges in order to reproduce the original analysis. Here, we averaged the DP of all within-network edges by their respective network, including edges with zero DP. Secondly, we repeated the analysis including between-network connections. We then averaged the DP between and within the different resting-state networks, creating a complete network-by-network DP matrix.

### Psychometric prediction

For prediction, we used the Connectome-based Predictive Modelling approach (CPM; Shen et al., 2017) and adapted the openly available script from: https://www.nitrc.org/frs/?group_id=51.

Using this framework, we predicted 30 psychometric variables supplied in the HCP dataset (see Supplements). In our main analysis, we focus on 3 behavioural variables of interest and provide further results in the supplement. Specifically, we focused on the fluid intelligence score assessed by Penn Progressive Matrices for its previous use by Finn et al. (2015) and others (Sui, Jiang, Bustillo, & Calhoun, 2020). In order to broaden the scope of our analysis and examine psychometric variables unrelated to fluid intelligence, we selected two additional psychometric variables, grip strength and language comprehension (assessed using the Picture Vocabulary Task) based on their low correlation with fluid intelligence (*r* = .02 and *r* = .20 respectively; Suppl. Table 1 for all correlations).

Behavioural prediction with CPM consists of three steps: feature selection, model building, and prediction. Features are selected by calculating the Pearson correlation between each edge in the training set and the psychometric variable. Edges are separated into correlated and anti-correlated edges and thresholded. Here, we tested: p < 0.05, 0.01, 0.005 and 0.001 following previous work by Finn et al., (2015). Next, the thresholded correlated and anti-correlated edges are summed up, resulting in two summary values (a positive set and a negative set). Positive and negative summary values are used as a predictor of the measured cognitive variable in two linear regressions using least squares estimates. In the last step, positive and negative summary values are calculated for every subject in the test set using the same features identified during the feature selection step. The summary network strengths are then used to predict the cognitive variable. A detailed description of CPM can be found in (Finn et al., 2015; Shen et al., 2017).

We used 10-fold cross validation (CV) repeated 100 times, resulting in 10×100 measures of accuracy. To evaluate model accuracy, we collected the predicted cognitive scores for each subject in each repeat of our CV (i.e., 100 predictions per subject), and averaged across all repeats, following Nostro et al. (2018). To evaluate the significance of the relationship between the predicted and the measured scores, we performed permutation testing with 1000 permutations. In each permutation, we correlated the averaged predicted scores with the measured cognitive scores found in the HCP data. The *p*-value (right-tailed) was calculated by dividing the permutations that exceeded the non-permuted correlation value by the number of permutations plus one.

Lastly, we used SVR to evaluate the overlap of discriminatory and predictive edges independent of CPM. To this end, we repeated the above prediction procedure but removed the feature selection and model building steps and instead used SVR for model building. SVR parameters were set at default values with no hyperparameter optimization (linear kernel, C = 0.75, lambda = 0.0035). We extracted the highly predictive edges using the SVR weights, which were thresholded using the same method applied to the PP edges (described below).

### Overlap between differential power, predictive power, and high variability

In order to perform our overlap analysis, we required a binarized matrix of edges with high discriminatory potential and a binarized matrix of edges with high predictive power. For discriminatory power, we thresholded the complete DP matrix, using the 99th percentile of edge DP, resulting in a sparse binary matrix of high DP and non-high DP edges. For predictive power, all edges selected in at least 80% of CV folds during prediction were used, resulting in binarized matrix (PP). In a final step, the overlap was calculated by overlaying the thresholded DP and PP matrices and calculating the intersection of positive values. To assess the overlap of fingerprinting and high-variability edges, we overlaid the DP matrix with a matrix of the functional connectivity standard deviation across participants, thresholded at the 99^th^ percentile. Other thresholds were also assessed (see Supplements).

Furthermore, we investigated the distribution of the standard deviation in all edges of the connectome, in edges with high DP, and in edges with high PP for the different psychometric predictions. To assess if edge-to-edge overlaps were statistically larger than would be expected by chance, we performed a permutation test with 1000 permutations. In each permutation, we calculated the intersection between fingerprinting DP edges and a degree-preserving random matrix (preserving degrees of PP or SD matrix) using the brain connectivity toolbox (Rubinov & Sporns, 2010).

### Topographical localization

To localize regions important to either fingerprinting or the prediction of psychometric variables, we calculated the node degree for each region by summing up the number of connected edges in the sparse DP matrix (in fingerprinting) and in the sparse PP matrices (in psychometric prediction). To compare the topographical organization found in fingerprinting and in prediction, we calculated the Spearman correlation between their node degrees and tested for significance of the topological overlap using spin permutation testing with 5000 permutations (Alexander-Bloch et al., 2018; Váša et al., 2018). Spin permutation allows for correlational analyses of cerebral topology while conserving spatial data properties such as non-independence amongst neighbouring parcels.

## Results

We first replicated the high fingerprinting accuracies and within-network overlap presented by Finn et al. (2015). Next, we investigated the overlap between features of interest in fingerprinting and prediction on edge-by-edge, network-by-network, and a large-scale topographical level. Here, we only present the results for fluid intelligence, language comprehension, and grip strength for the positive models of the CPM framework, with a feature selection threshold of *p* < .01. However, results equivalently hold for the negative models as well as for other feature selection thresholds (p < .001, .005, .05), and can be found in the supplements (Suppl. Fig. 1-3).

### Within-network connections analysis

#### Connectome fingerprinting and network distribution

We observed high fingerprinting accuracy of 96.8% (328/339, permutation-derived *p*<.001 against chance) when identifying individuals from session 1, and 97.3% (330/339, p<0.001) when identifying individuals from session 2. Focussing on within-network connections, we observed strong involvement of highly discriminatory edges from the medial frontal (MFN), frontoparietal (FPN), and default mode network (DMN) as well as minor involvement of the subcortical-cerebellar network (SCN) (Fig. 1a). These results closely resemble findings by Finn et al. (2015), who found comparable accuracies and within-network contributions.

**Figure 1.**
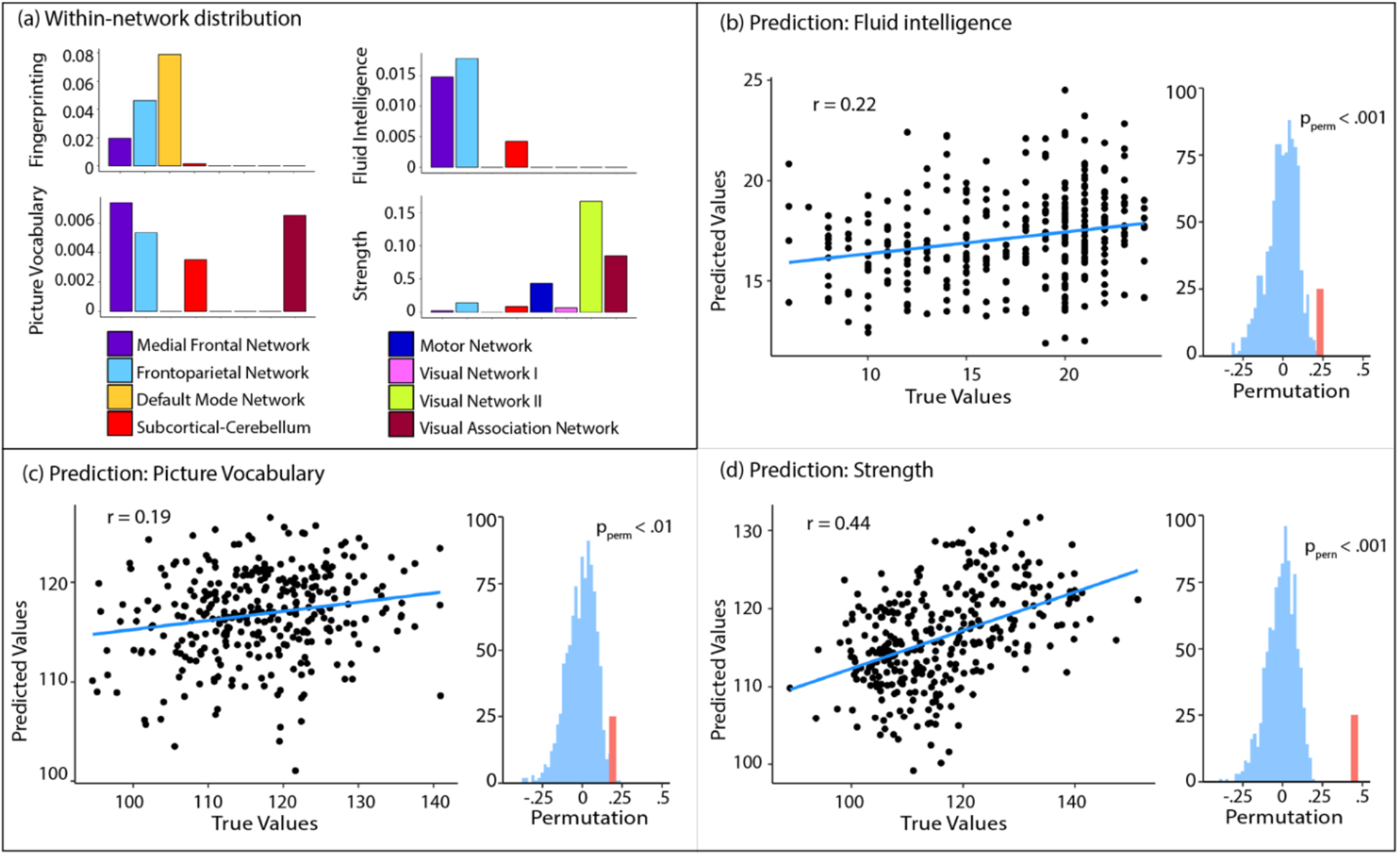
Within-network distribution of selected edges and behavioural prediction. Panel (a) visualizes the percentage of selected edges for fingerprinting and for prediction within each network, adjusted for the total number of edges in each network. Panels (b-d) show the prediction results of three psychometric variables of interest. Language comprehension was evaluated using the picture vocabulary task.

#### Behavioural prediction

We found a significant correlation between the measured values and the predicted values of fluid intelligence (r = .22, *p* < .001; Fig. 1b). The within-network connections that were most often selected as features were found in the MFN, FPN and SCN (Fig. 1a). Akin to Finn et al. (2015), networks supporting prediction resembled the networks displaying the highest proportion of discriminatory edges in fingerprinting, i.e. both the MFN and FPN contributed to fingerprinting and prediction of fluid intelligence (Fig 1a). Further corroborating these findings, we found the DMN to be involved in the individual fingerprints but not in the positive prediction models, as was reported in (Finn et al., 2015). Motor Network (MN), Visual Network I (VN1), Visual Network II (VN2) and the Visual Association Network (VASN) did not strongly contribute to prediction or fingerprinting, as edges from those networks did not appear in the 99^th^ percentile of discriminatory nor predictive edges.

Overall, we observed a significant correlation between measured and predicted values in 12 out of 30 psychometric variables (Suppl. Table 2 for all prediction results) including our two other variables of interest, language comprehension and grip strength (Table 1, all p-values permutation-derived, with n = 1000). When we examined the network contributions to the prediction of language comprehension, we found large involvement of within-network edges in the MFN, FPN, and VASN, and more discrete involvement of the SCN within-network edges. Once more, these networks resembled those best discriminating between individuals. In strength prediction, VN2 and the VASN within-network edges were most predictive. In sum, within-network analyses of discriminatory and predictive edges seem to suggest that connectome fingerprints may be relevant to inter-individual differences in higher-order cognitive functions such as fluid intelligence and language comprehension, but not grip strength, in line with previous reports (Finn et al., 2015).

### Overlap analysis

The above findings notwithstanding, if participant identification and behavioural prediction truly rest on the same functional connectome signatures, significant overlap between discriminatory and predictive features would be expected beyond the mere resemblance of within-network contributions, i.e., at the level of single edges, between-network connections, and the large-scale spatial distribution of discriminatory and predictive nodes.

#### Single-edge analysis

First, we investigated the overlap between highly discriminatory edges in fingerprinting and edges predictive of behaviour, without averaging or grouping these edges into resting-state networks (Table 2 and Fig. 2a). We found that the overlap between discriminatory and predictive edges did not exceed chance level for any of the psychometric variables (Table 2).

**Table 2.**
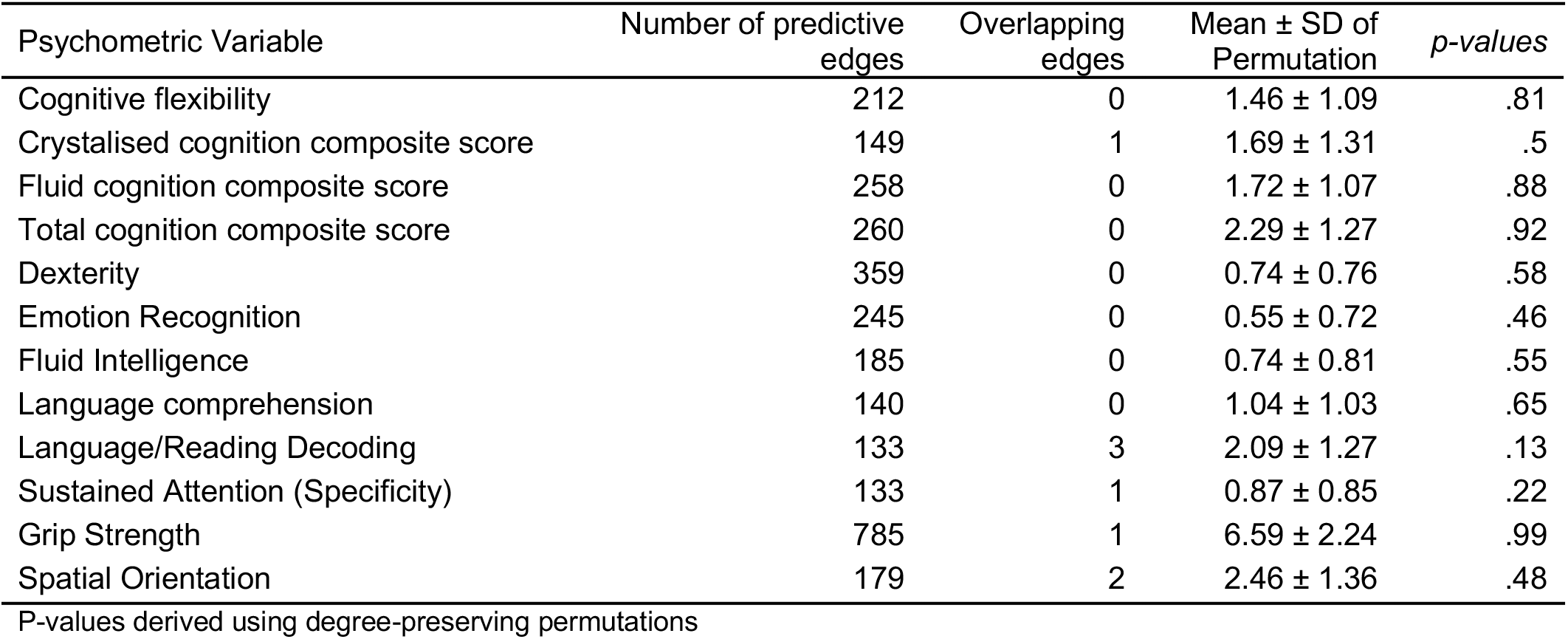
Overlap between edges with high predictive power and fingerprints

| Psychometric Variable | Number of predictive edges | Overlapping edges | Mean ± SD of Permutation | *p-values* |
| --- | --- | --- | --- | --- |
| Cognitive flexibility | 212 | 0 | 1.46 ± 1.09 | .81 |
| Crystallised cognition composite score | 149 | 1 | 1.69 ± 1.31 | .5 |
| Fluid cognition composite score | 258 | 0 | 1.72 ± 1.07 | .88 |
| Total cognition composite score | 260 | 0 | 2.29 ± 1.27 | .92 |
| Dexterity | 359 | 0 | 0.74 ± 0.76 | .58 |
| Emotion Recognition | 245 | 0 | 0.55 ± 0.72 | .46 |
| Fluid Intelligence | 185 | 0 | 0.74 ± 0.81 | .55 |
| Language comprehension | 140 | 0 | 1.04 ± 1.03 | .65 |
| Language/Reading Decoding | 133 | 3 | 2.09 ± 1.27 | .13 |
| Sustained Attention (Specificity) | 133 | 1 | 0.87 ± 0.85 | .22 |
| Grip Strength | 785 | 1 | 6.59 ± 2.24 | .99 |
| Spatial Orientation | 179 | 2 | 2.46 ± 1.36 | .48 |

P-values derived using degree-preserving permutations

**Figure 2.**
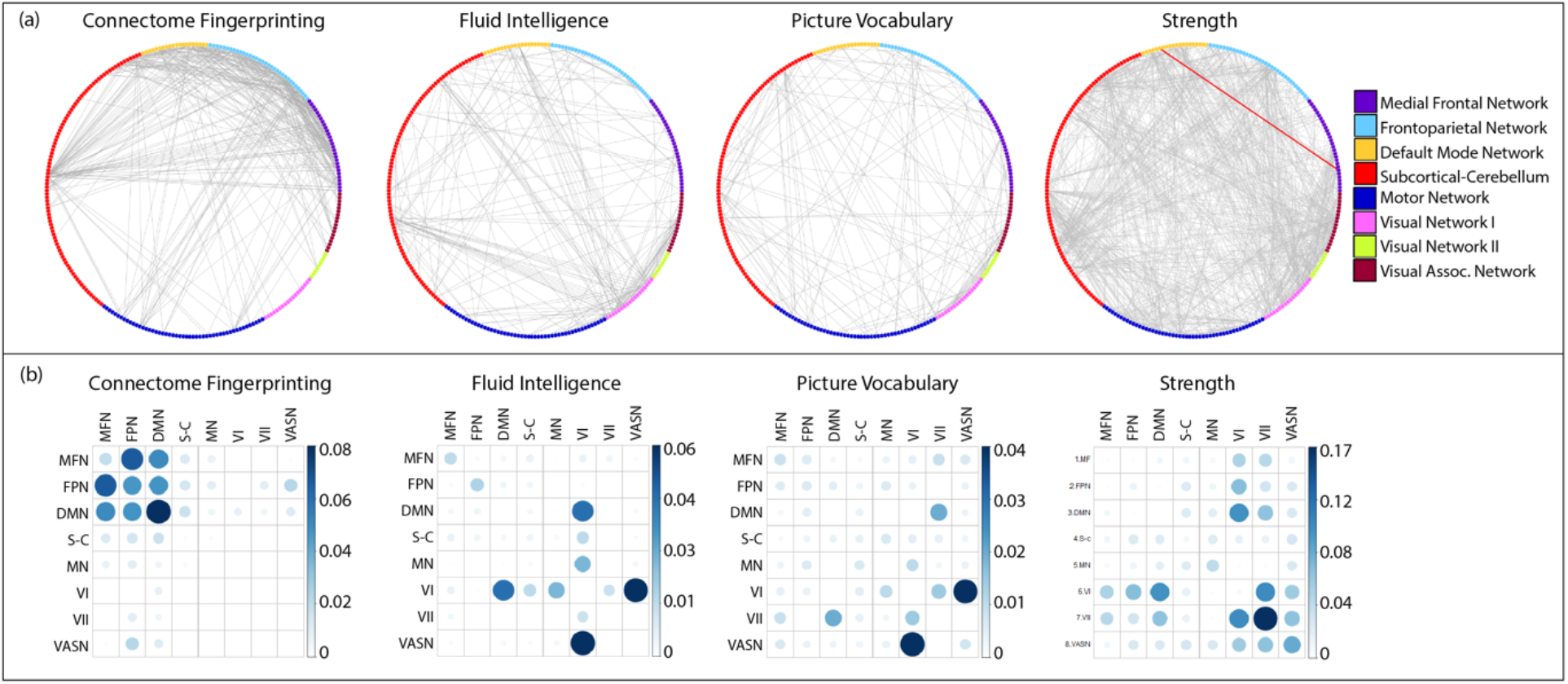
Single-edge and between-network overlap for fingerprints and prediction. In panel (a), grey lines mark highly discriminatory edges for fingerprints and predictive edges for behaviour, both thresholded at the 99^th^ percentile. Red lines, only available for strength, visualize overlap. For fluid intelligence and language comprehension, no edges overlapped. Panel (b) shows the entire network-by-network matrix of selected edges, adjusted for total number of edges.

Furthermore, if discriminatory edges are indeed relevant to the psychometric variables of interest, one should be able to predict these scores using the discriminatory edges directly. We tested this by modifying our prediction pipeline, replacing the feature selection step and instead directly applying the fingerprinting edges in the training data of each CV fold. We then used this set of discriminatory edges to predict fluid intelligence, language comprehension, and grip strength in the test set. We found that predictions based on discriminatory edges could not significantly predict any of the three behavioural variables (Table 3), further corroborating that individual fingerprints are not related to behaviour on the single-edge level.

**Table 3.**
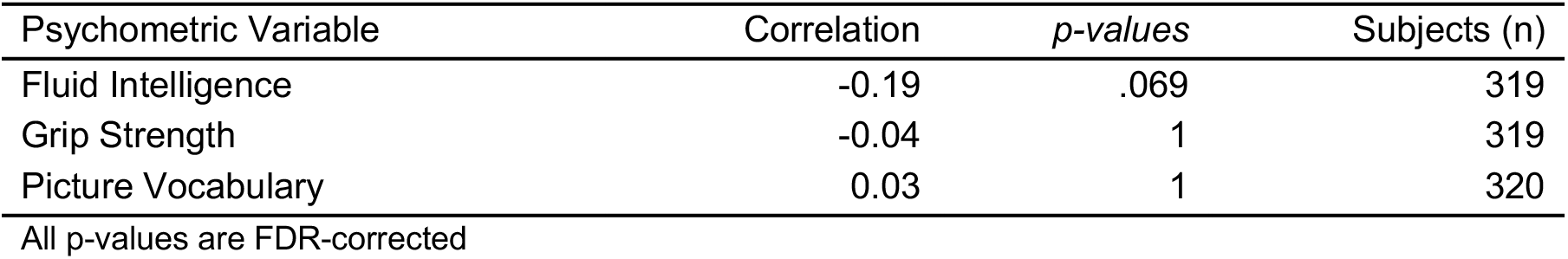
Behavioural prediction results using fingerprinting edges for model construction

#### Network analysis

Next, as individual functional connection weights have low reliability (Noble, Scheinost, & Constable, 2019), we investigated the network distribution of edges, this time including both within- and between-network connections (Fig. 2b). For fingerprinting, we found a cluster of highly connected edges between as well as within the MFN, FPN, and DMN, and to a lesser extent in the SCN. This was in stark contrast to between-network connections found in psychometric prediction, which displayed a much more variable pattern. Most reliably, predictive features included connections between the DMN, the visual networks, and the rest of the brain, with the exception of some within-network edges in the MFN and FPN for fluid intelligence. Furthermore, analysing the proportion of selected edges by network, we found that fingerprints did not significantly relate to fluid intelligence (*r* = −.08, *p* = .620), language comprehension (*r* = −.15, *p* = 0.585), nor grip strength (*r* = −.39, *p* = .051). Taken together, these findings suggest that even on a network level, individual fingerprints were not related to behaviour.

#### Topological analysis of nodes with high degree of predictive edges

Next, we investigated the overlap on a large-scale topological level. We found that discriminatory nodes (i.e., nodes with high degree of discriminatory edges) clustered almost exclusively in the superior frontal, inferior parietal, and superior temporal regions (Fig. 3a). In line with our network-derived findings, predictive nodes displayed a more variable spatial distribution (Fig. 3b-d) and, importantly, largely covered different parts of the cortex compared to discriminatory nodes. Corroborating these observations, there was no correlation between the spatial distribution of nodes in fingerprinting and any behavioural prediction (Fig. 3b-d, right panel, all p-values derived using spin permutation), again suggesting the spatial organisation of discriminatory nodes was not related to behaviour.

**Figure 3.**
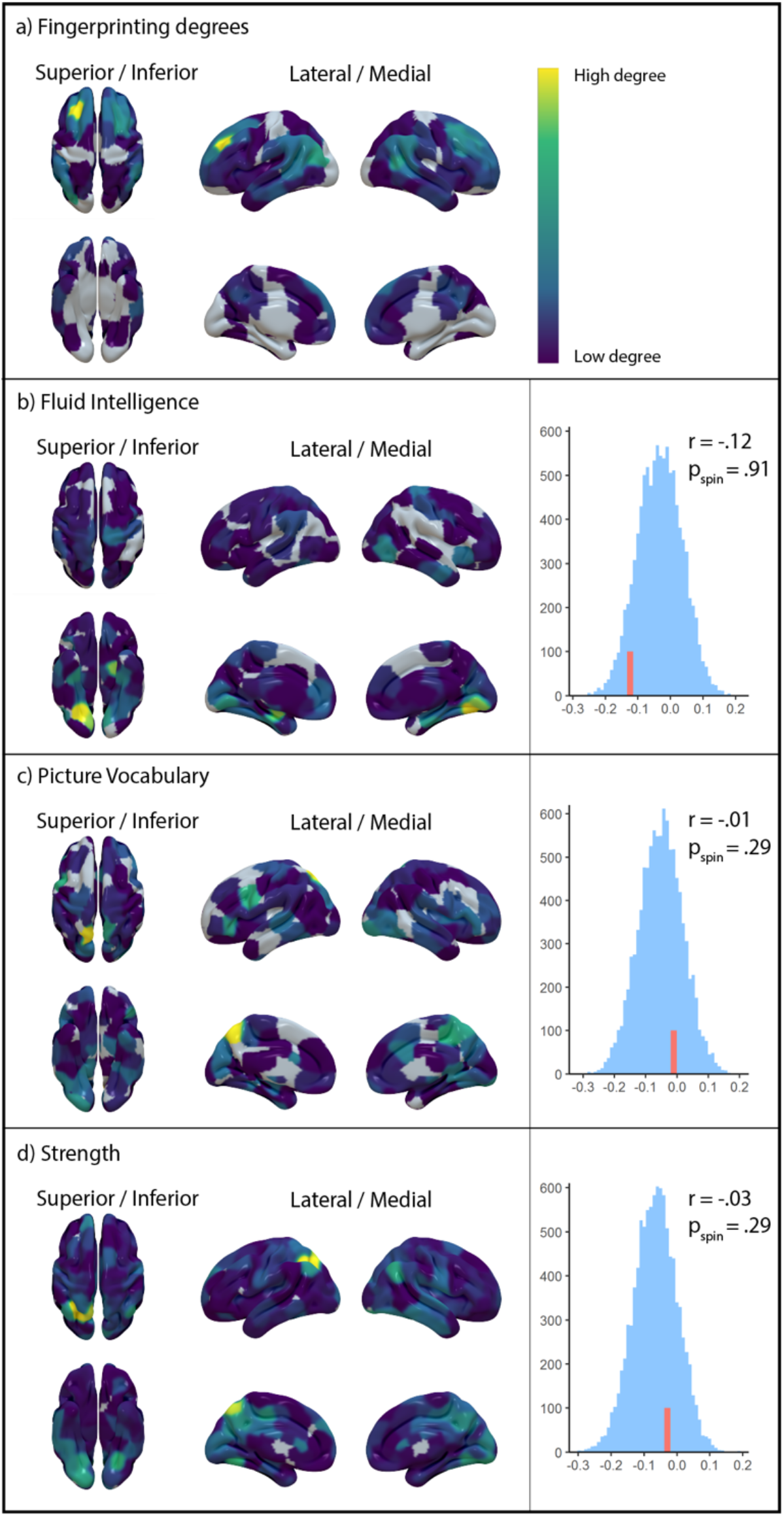
Spatial distribution of node degrees. Distribution of node degrees for discriminatory nodes (a) and behaviourally predictive nodes (b-d) on the left. The edges underlying the node degrees are thresholded at the 99^th^ percentile of discriminatory potential or predictive power. The right-hand side displays the spin permutation results, with red lines marking the empirical correlation of discriminatory and predictive nodes.

### Variability analysis

Subsequently, we performed an exploratory analysis to investigate how edge properties relate to the divergence between discriminatory and predictive connectome features. Focussing on edge standard deviation of functional connectomes, we discovered that discriminatory edges generally show high variability across participants (Fig. 4). Investigating the variability by comparing the 99^th^ percentile of discriminatory edges in fingerprinting and the 99^th^ percentile of edges with high standard deviation showed a strong and significant (122/358, *p* < .001) overlap, which increased even further when thresholding at the 98^th^ percentile (179/358, *p* < .001) and the 95^th^ percentile (286/358, *p* < .001; all *p*-values permutation-derived, Fig. 4a). Furthermore, edges used in prediction showed significantly lower variability for all three psychometric variables (*p* < .001, FDR-corrected) than discriminatory edges (Fig. 4b). Taken together, fingerprinting signatures significantly overlap with edges showing higher variability in connectivity across subjects, while edges predictive of behaviour are constrained to edges with intermediate variability.

**Figure 4.**
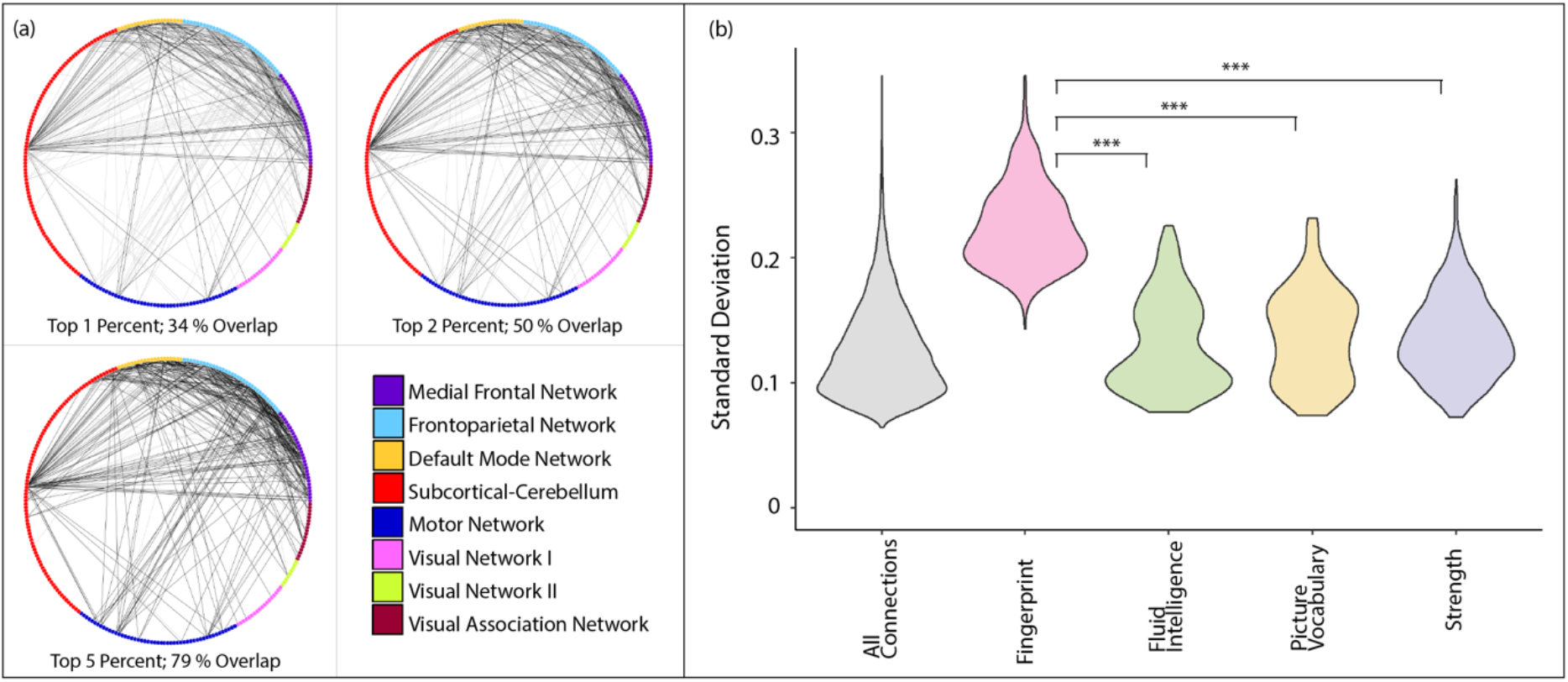
Overlap of fingerprints and high-variability edges. In panel (a), black lines designate overlapping edges between the top one, two or five percent of discriminatory edges and high-variability edges. Grey lines depict lower-variability discriminatory edges. Panel (b) shows the distribution of edge standard deviation across participants. *** p < .001, FDR-corrected.

### Validation analysis

Firstly, we repeated the analysis of edge-level overlap between discriminatory edges and predictive edges for all three behaviors, using Support Vector Regression instead of CPM. This independent prediction method corroborated the lack of overlap for all tested behaviors (Suppl. Table 3). Additionally, we tested whether we could replicate our findings using different parcellation schemes. Focusing on the prediction of fluid intelligence, we observed significant correlations between predicted and measured intelligence scores using CPM with all three atlases (Brainnetome: *r* = .26, *p* < .001, HCP: *r* = .18, *p* = .001, AAL: *r* = .19, *p* < .001). We also achieved fingerprinting accuracies of more than 90% for all atlases, with the HCP MMP1 atlas resulting in accuracies of up to 99% (Suppl. Table 4). Our findings concerning a lack of overlap between discriminatory and predictive edges held true for between-network, anatomical and single-edge overlap (Brainnetome: n = 3/301, *p* = .137, HCP: n = 3/646, *p* = .164, AAL: n = 0/67, *p* = .269) in all three parcellation schemes (Fig. 5a-b). We were also able to replicate the relationship between edge-variability and fingerprinting, showing a high overlap between the most discriminatory edges and edges with high standard deviation (Fig. 5c), as well as the significantly lower variability in edges predictive of behaviour (Fig. 5d).

**Figure 5.**
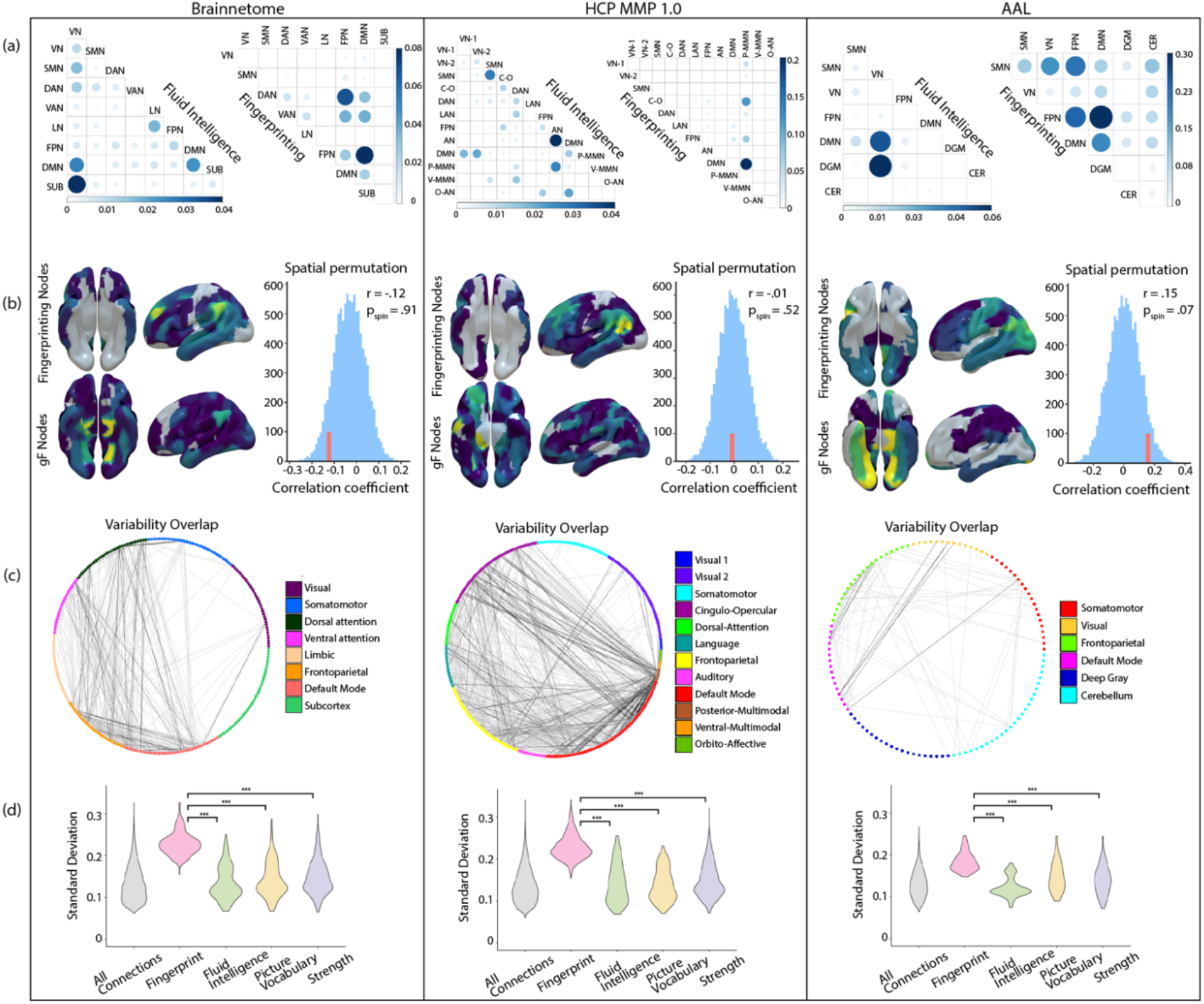
Control analyses for different functional atlases. Results for Brainnetome, HCP MMP 1.0, and AAL atlases (left to right). From top to bottom, panels visualize (a) within and across-network connections, (b) spatial topology and spin permutation of nodes for fluid intelligence and fingerprinting, (c) the overlap of individual edges between fingerprints and high-variability edges, and (d) the distribution of edge standard deviations over participants in fingerprinting and behavioural prediction. *** p < .001, FDR-corrected.

## Discussion

In the present study, we show that fingerprinting signatures and behavioural prediction rest on highly distinct functional systems of the human connectome. We were able to replicate the seminal findings by Finn et at. (2015) demonstrating high accuracy in participant identification with connectome fingerprinting as well as the importance of within-network edges in higher-order resting-state networks for both prediction and fingerprinting. These findings have been interpreted as supporting the functional relevance of fingerprinting signatures, that is, networks that best discriminate individuals from one another are also strongly involved in cognitive function (Finn et al., 2015, 2017). However, these findings were restricted to a specific level of analysis (group-level within-network connections), necessitating further exploration into whether participant identification and behavioural prediction truly rest on the same functional connectome signatures.

Here, we found evidence to the contrary, as there was a strong divergence between functional signatures supporting prediction of behaviour and connectome fingerprints. This held true on a network level, if we considered both within as well as between-network connections, on the level of single edges, and on the level of large-scale spatial organisation of discriminatory and predictive nodes. Thus, comparing the underlying patterns recruited during prediction and fingerprinting, we found no overlap that could justify the argued relationship between them. Additionally, as a positive control, we directly used individual fingerprints for the prediction of behaviour and found this to be unsuccessful, further corroborating our findings. To address the many degrees of freedom in the design of the analysis, we also show that our findings are highly robust against varying methodological choices. Specifically, we used four parcellation schemes, two prediction methods, and tested different feature selection thresholds. In sum, the results presented here suggest that discriminatory and predictive signatures of the human connectome rely on highly distinct functional systems.

In this regard, our findings expand on the recent notion of a dichotomy between resources valuable to subject identification and behavioural prediction. While this dichotomy has been shown to apply with respect to different imaging modalities (Mansour L, Tian, Yeo, Cropley, & Zalesky, 2021) here we show the separation to be present within a single modality. In this context, we here discovered that the variability of individual edges strongly distinguishes between fingerprinting and prediction and we propose this variability to be at the root of the dichotomy. Discriminatory edges, but not predictive ones, showed a substantial overlap with edges that are highly variable across participants. The different mechanisms underlying fingerprinting and prediction might clarify these findings. As an example, CPM selects edges which show significant correlation between functional connectivity and a behavioural outcome measure in the sample during feature selection. Thus, CPM is still a group-level procedure and requires edge variation to be linearly related to behaviour in order to be selected for prediction. In contrast, in fingerprinting, edges are selected based on intra-subject similarity, given sufficient inter-subject variability. Importantly, here, no group-relationship is considered. CPM and fingerprinting thus relate differently to edge variability, i.e., edges selected in CPM need to covary with behaviour, whereas fingerprinting is impartial to the source of edge variability. As a consequence, the high variability of functional connections selected in fingerprinting could result from a range of sources. For example, variations could stem from differences in functional network topology (Gordon, Laumann, Adeyemo, & Petersen, 2017) or structural variability such as differences in cortical thickness (Mueller et al., 2013), or folding patterns (Duan et al., 2019). These differences might also result in stable variation in functional connectivity, whilst not necessarily relating to behaviour in a linear fashion. We believe that the increased variability in multimodal brain regions (Mueller et al., 2013; Paquola et al., 2019; Seitzman et al., 2019) leads to a higher likelihood of individual variants from different sources. As a consequence, we observe clusters of discriminatory edges in these regions when averaging the discriminatory potential of individual edges over all participants. The significance of these structural and functional variations is difficult to discern, our results point to the variation exploited during fingerprinting not being related to behaviour. Further research will be necessary to establish whether edge variability also serves as a separating marker in other imaging modalities and whether the findings by (Mansour L et al., 2021) might also be supported by the proposed relationship between fingerprinting, behavioural prediction, and signal variability.

### Limitations

We closely followed Finn et al. (2015) in the pre-processing steps and used the same methods for the identification of subjects, the prediction of behaviour and the extraction of high-value edges for fingerprinting (Shen et al., 2017). Furthermore, we mirrored the data analysis pipeline, initially focussing on within-network edges. Nonetheless, while we found functional connectivity to be a significant predictor of fluid intelligence with an accuracy similar to other published work (Ferguson, Anderson, & Spreng, 2017; Greene, Gao, Scheinost, & Constable, 2018), we did not achieve the high prediction scores reported in Finn et al. (2015). There are different possible explanations for this, one of them being our use of the unrelated sample from the HCP database. This sample has the advantage of being larger and thus more robust to overfitting, and it assures the independence between subjects during cross-validation (Poldrack, Huckins, & Varoquaux, 2020). However, this independence might have influenced our prediction accuracies. Furthermore, we used 10-fold cross validation instead of leave-one-out cross validation (Varoquaux et al., 2017).

## Conclusion

In contrast to initial reports linking connectome fingerprinting signatures directly to behaviour, we here show in detail that participant identification and behavioural prediction from individual connectomes rely on highly distinct functional brain systems. This divergence raises the question what the variability sustaining individual fingerprints ultimately relates to. Parsimony suggests that neurological variation should also be linked to phenotypic presentation, yet our results indicate that there is no simple one-to-one mapping between function and fingerprints. As such, further methodological development and conceptualization will be necessary to deepen our understanding of individual functional signatures and their behavioural and biological significance.

