## Supplemental Material for "Fingerprinting and behavioural prediction rest on distinct functional systems of the human connectome"

SUPPLEMENT FIGURES AND TABLES

**Supplementary Table 1:** Correlations between all 30 behavioral measures

|  | FCCS | CCCS | TCCS | CF | gF | SA | GS | D | L/RD | LC | SO | ER |
| --- | --- | --- | --- | --- | --- | --- | --- | --- | --- | --- | --- | --- |
| Fluid cognition composite | 1 | 0.21 | 0.94 | 0.7 | 0.16 | 0.04 | 0.06 | 0.11 | 0.22 | 0.16 | 0.16 | 0.1 |
| Crystalised cognition composite | 0.21 | 1 | 0.51 | 0.17 | 0.21 | -0.06 | 0.03 | 0.22 | 0.93 | 0.9 | 0.16 | -0.01 |
| Total cognition composite | 0.94 | 0.51 | 1 | 0.66 | 0.2 | 0.01 | 0.06 | 0.15 | 0.49 | 0.44 | 0.18 | 0.07 |
| Cognitive flexibility | 0.7 | 0.17 | 0.66 | 1 | 0.16 | 0.06 | 0.08 | 0.08 | 0.2 | 0.09 | 0.17 | 0.1 |
| Fluid Intelligence | 0.16 | 0.21 | 0.2 | 0.16 | 1 | 0.81 | 0.02 | 0.13 | 0.19 | 0.2 | 0.8 | 0.8 |
| Sustained Attention | 0.04 | -0.06 | 0.01 | 0.06 | 0.81 | 1 | 0.02 | 0.03 | -0.06 | -0.04 | 0.83 | 0.94 |
| Grip strength | 0.06 | 0.03 | 0.06 | 0.08 | 0.02 | 0.02 | 1 | -0.13 | 0.05 | -0.01 | 0.03 | 0.03 |
| Dexterity | 0.11 | 0.22 | 0.15 | 0.08 | 0.13 | 0.03 | -0.13 | 1 | 0.2 | 0.2 | 0.07 | 0.08 |
| Language/Reading Decoding | 0.22 | 0.93 | 0.49 | 0.2 | 0.19 | -0.06 | 0.05 | 0.2 | 1 | 0.67 | 0.15 | -0.03 |
| Language comprehension | 0.16 | 0.9 | 0.44 | 0.09 | 0.2 | -0.04 | -0.01 | 0.2 | 0.67 | 1 | 0.14 | 0.01 |
| Spatial Orientation | 0.16 | 0.16 | 0.18 | 0.17 | 0.8 | 0.83 | 0.03 | 0.07 | 0.15 | 0.14 | 1 | 0.81 |
| Emotion Recognition | 0.1 | -0.01 | 0.07 | 0.1 | 0.8 | 0.94 | 0.03 | 0.08 | -0.03 | 0.01 | 0.81 | 1 |

**Supplementary Table 2:** Results for all 30 behavioral predictions for positive and negative models

| Behaviour | Correlation  Positive model | *p value* | MSE positive model | Edges  Positive model | Correlation  Negative Model | *p value* | MSE negative model | Edges  Negative model | Subjects (n) | Adjusted *p* positive | Adjusted *p* negative |
| --- | --- | --- | --- | --- | --- | --- | --- | --- | --- | --- | --- |
| Fluid cognition composite | 0.21 | < .001 | 136.97 | 1298 | 0.25 | < .001 | 133.74 | 1242 | 318 | .002 | < .001 |
| Crystalised cognition composite | 0.21 | < .001 | 100.82 | 754 | 0.16 | .005 | 105.53 | 648 | 320 | .001 | .028 |
| Total cognition composite | 0.25 | < .001 | 208.57 | 1198 | 0.27 | < .001 | 203.76 | 1162 | 318 | < .001 | < .001 |
| MMSE | -0.06 | .277 | 1.38 | 312 | -0.01 | .842 | 1.34 | 304 | 320 | 0.79 | 1 |
| Cognitive flexibility | 0.18 | .001 | 104.89 | 1522 | 0.21 | < .001 | 103.61 | 1746 | 319 | .006 | .002 |
| Inhibition | 0.12 | .028 | 111.43 | 678 | 0.06 | .318 | 118.18 | 538 | 320 | .12 | 1 |
| Fluid Intelligence | 0.22 | < .001 | 22.4 | 798 | 0.15 | .008 | 23.72 | 528 | 319 | < .001 | .045 |
| Processing Speed | 0.14 | .015 | 261.23 | 896 | 0.18 | .001 | 252.13 | 1010 | 320 | .071 | .01 |
| Working Memory | 0.09 | .102 | 136.68 | 730 | 0.16 | .004 | 129.47 | 864 | 320 | .307 | .022 |
| Sustained Attention (Sensitivity) | -0.01 | .855 | 0.004 | 276 | -0.02 | .783 | 0.004 | 232 | 319 | 1 | 1 |
| Sustained Attention (Specificity) | 0.17 | .003 | 0.001 | 636 | 0.12 | .026 | 0.001 | 692 | 319 | .015 | .105 |
| Personality: Agreeableness | 0.1 | .077 | 38.83 | 520 | 0.07 | .236 | 39.26 | 556 | 319 | .256 | .786 |
| Personality: Openness | 0.01 | .893 | 44.46 | 382 | -0.04 | .51 | 45.62 | 262 | 319 | 1 | 1 |
| Personality: Conscientiousness | 0.03 | .584 | 35.58 | 332 | -0.04 | .466 | 37.14 | 374 | 319 | 1 | 1 |
| Personality: Neuroticism | 0.05 | .392 | 63.47 | 392 | 0.07 | .206 | 61.98 | 496 | 319 | .979 | .728 |
| Personality: Extraversion | 0.05 | .357 | 41.11 | 348 | 0 | .978 | 42.68 | 278 | 319 | .931 | 1 |
| Grip strength | 0.44 | < .001 | 101.09 | 2534 | 0.42 | < .001 | 104.07 | 2106 | 319 | < .001 | < .001 |
| Dexterity | 0.23 | < .001 | 121.06 | 1230 | 0.21 | < .001 | 124.96 | 1176 | 320 | < .001 | .002 |
| Audition | 0.06 | .306 | 2.42 | 348 | 0.05 | .357 | 2.46 | 364 | 317 | .836 | 1 |
| Smell | -0.12 | .038 | 99.58 | 186 | 0.02 | .714 | 92.96 | 296 | 319 | .153 | 1 |
| Taste | 0.03 | .637 | 236 | 532 | 0.12 | .034 | 214.94 | 526 | 318 | 1 | .126 |
| Sleep Quality | 0.03 | .606 | 9.71 | 494 | 0.03 | .628 | 9.76 | 556 | 320 | 1 | 1 |
| Episodic Memory | 0.09 | .104 | 189.55 | 476 | 0.17 | .003 | 181.06 | 436 | 320 | .307 | .019 |
| Language/Reading Decoding | 0.19 | < .001 | 126.58 | 632 | 0.14 | .01 | 131.24 | 418 | 320 | .003 | .047 |
| Language comprehension | 0.19 | < .001 | 88.69 | 752 | 0.17 | .002 | 89.95 | 618 | 320 | .004 | .014 |
| Delay Discounting 200 | 0.11 | .053 | 0.05 | 524 | 0.03 | .621 | 0.06 | 388 | 319 | .198 | 1 |
| Delay Discounting 40K | 0.02 | .692 | 0.1 | 412 | 0.05 | .399 | 0.1 | 360 | 319 | 1 | 1 |
| Spatial Orientation | 0.21 | < .001 | 19.11 | 914 | 0.16 | .003 | 19.69 | 770 | 319 | .002 | .021 |
| Emotion Recognition | 0.2 | < .001 | 7.08 | 1112 | 0.13 | .02 | 7.56 | 942 | 319 | .002 | .086 |
| Emotion Recognition (hits) | 0.1 | .062 | 156502.47 | 468 | 0 | .995 | 168863.67 | 216 | 319 | .218 | 1 |

**Supplementary Table 3:** SVR prediction and overlap analysis

| Psychometric Variable | Prediction (r value); *p-value* | Predictive edges (n) | Edges Overlapping (n) with fingerprints | Mean ± SD of Permutation | *p-values* |
| --- | --- | --- | --- | --- | --- |
| Fluid Intelligence | .29; <.001 | 147 | 7 | 5.38 ± 2.14 | .151 |
| Language Comprehension | .34; <.001 | 155 | 6 | 5.54 ± 2.19 | .310 |
| Strength | .55; <.001 | 184 | 4 | 4.82 ± 2.02 | .555 |

**Supplementary Table 4:** Fingerprinting accuracies in validation atlases

| Parcellation Scheme | Percentage hits (R1 -> R2) | Percentage hits (R2-> R1) |
| --- | --- | --- |
| Shen (main text) | 96.7 | 97.3 |
| HCP MMP 1.0 | 99.1 | 99,1 |
| Brainnetome | 96.8 | 97.3 |
| AAL | 90.9 | 91.2 |

**Supplementary Table 5:** Overlap analysis for all thresholds

| Behaviour | Threshold | Model | N_edges  Fingerprint | N_edges  Prediction | Overlap | Mean overlap  Permutation | SD overlap  Permutation | *P values* |
| --- | --- | --- | --- | --- | --- | --- | --- | --- |
| Fluid Intelligence | .001 | neg | 36 | 10 | 0 | 0 | 0 | 1 |
| Fluid Intelligence | .001 | pos | 36 | 20 | 0 | 0 | 0 | 1 |
| Language Comprehension | .001 | neg | 36 | 12 | 0 | 0 | 0 | 1 |
| Language Comprehension | .001 | pos | 36 | 14 | 0 | 0 | 0 | 1 |
| Strength | .001 | neg | 36 | 165 | 0 | 0.01 | 0.09 | 1 |
| Strength | .001 | pos | 36 | 221 | 0 | 0.06 | 0.23 | 1 |
| Fluid Intelligence | .005 | neg | 179 | 48 | 0 | 0.59 | 0.71 | 1 |
| Fluid Intelligence | .005 | pos | 179 | 93 | 0 | 0.21 | 0.46 | 1 |
| Language Comprehension | .005 | neg | 179 | 51 | 0 | 0.56 | 0.69 | 1 |
| Language Comprehension | .005 | pos | 179 | 61 | 0 | 0.25 | 0.5 | 1 |
| Strength | .005 | neg | 179 | 413 | 2 | 0.43 | 0.64 | .496 |
| Strength | .005 | pos | 179 | 531 | 1 | 1.15 | 1.02 | 1 |
| Fluid Intelligence | .01 | neg | 358 | 109 | 1 | 2.58 | 1.36 | 1 |
| Fluid Intelligence | .01 | pos | 358 | 185 | 0 | 0.71 | 0.82 | 1 |
| Language Comprehension | .01 | neg | 358 | 103 | 1 | 1.35 | 1.11 | 1 |
| Language Comprehension | .01 | pos | 358 | 140 | 0 | 1.2 | 1.04 | 1 |
| Strength | .01 | neg | 358 | 667 | 8 | 3.33 | 1.73 | .132 |
| Strength | .01 | pos | 358 | 785 | 1 | 6.65 | 2.3 | 1 |
| Fluid Intelligence | .05 | neg | 1789 | 552 | 24 | 39.63 | 5.34 | 1 |
| Fluid Intelligence | .05 | pos | 1789 | 817 | 39 | 25.92 | 4.37 | .072 |
| Language Comprehension | .05 | neg | 1789 | 568 | 41 | 42.31 | 5.27 | 1 |
| Language Comprehension | .05 | pos | 1789 | 717 | 32 | 41.46 | 5.34 | 1 |
| Strength | .05 | neg | 1789 | 1738 | 83 | 77.84 | 7.63 | 1 |
| Strength | .05 | pos | 1789 | 2106 | 113 | 107.58 | 8.35 | 1 |

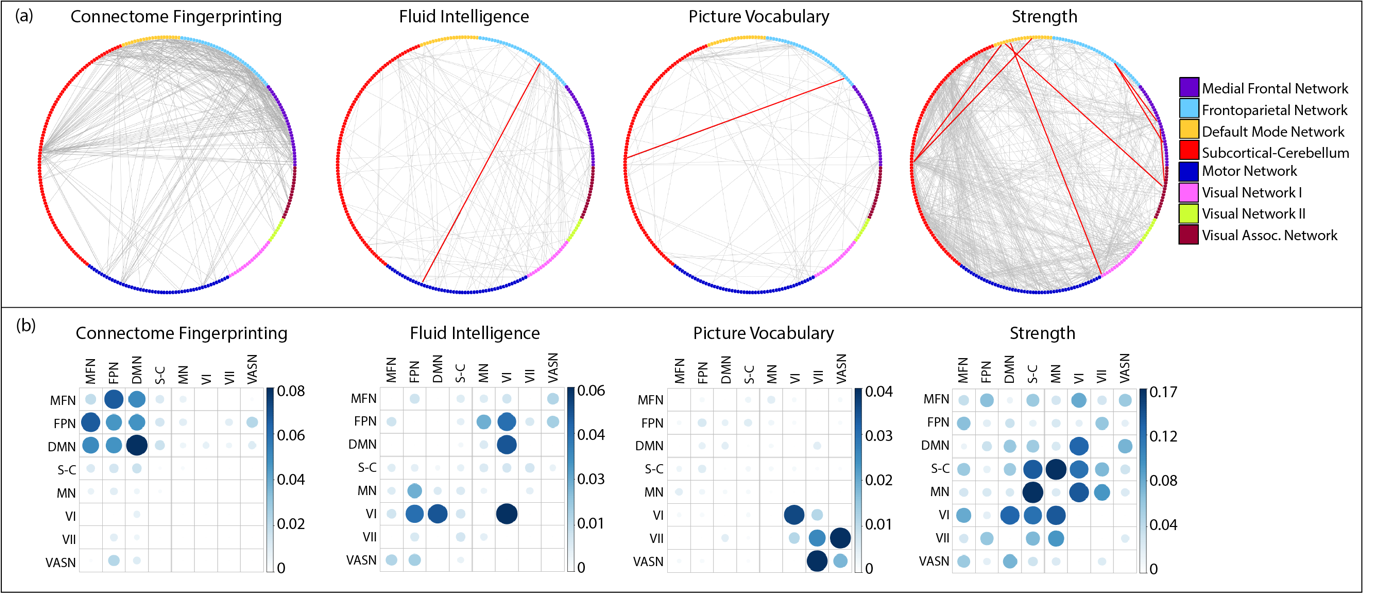

**Supplementary Figure 1**: Single-edge and between-network overlap for fingerprints and the negative prediction models, all thresholded at the 99^th^ percentile.

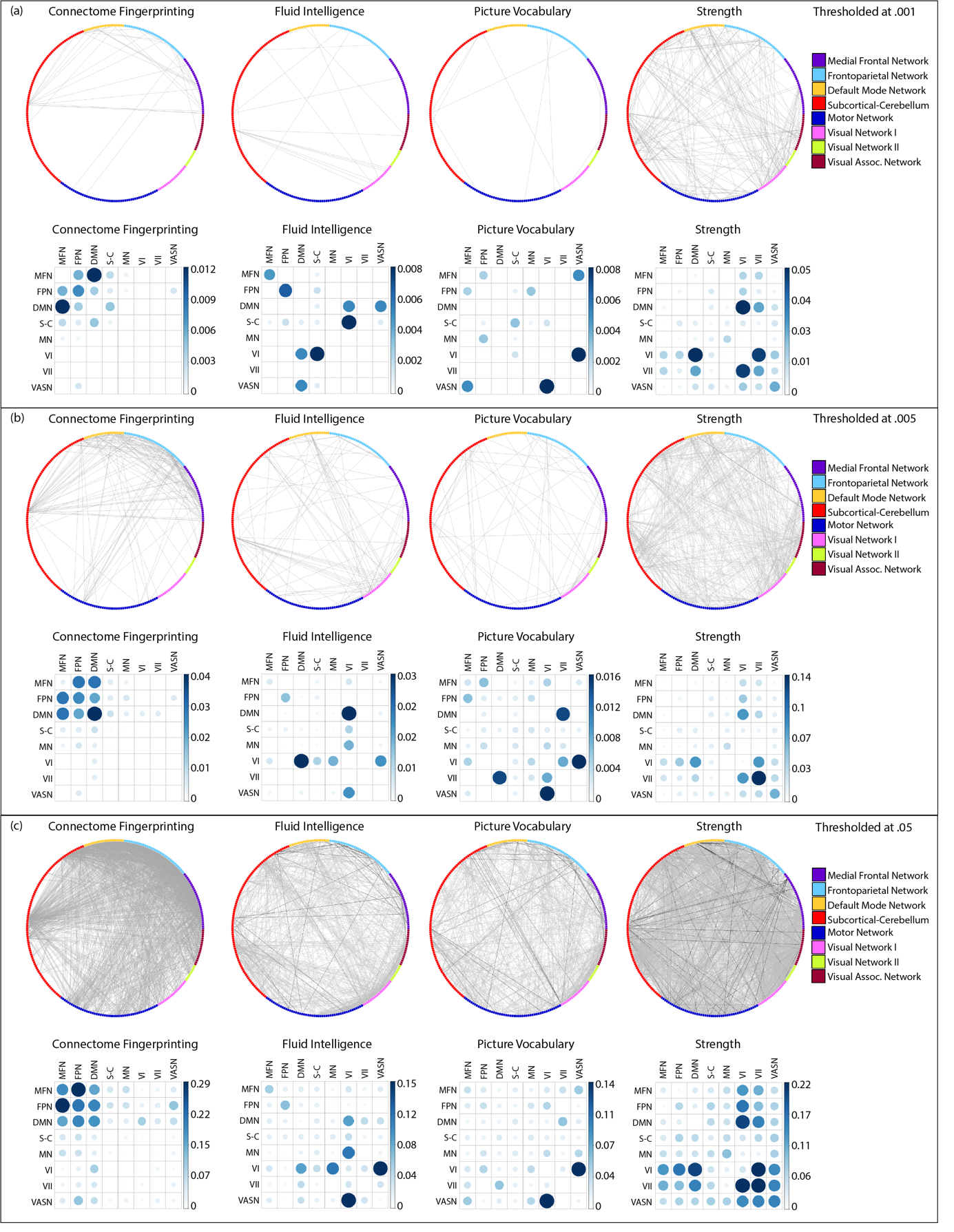

**Supplementary Figure 2:** Single-edge and between-network overlap for fingerprints and the positive prediction models, thresholded at the 99.9^th^, 99.5^th^ and 95^th^ percentile.

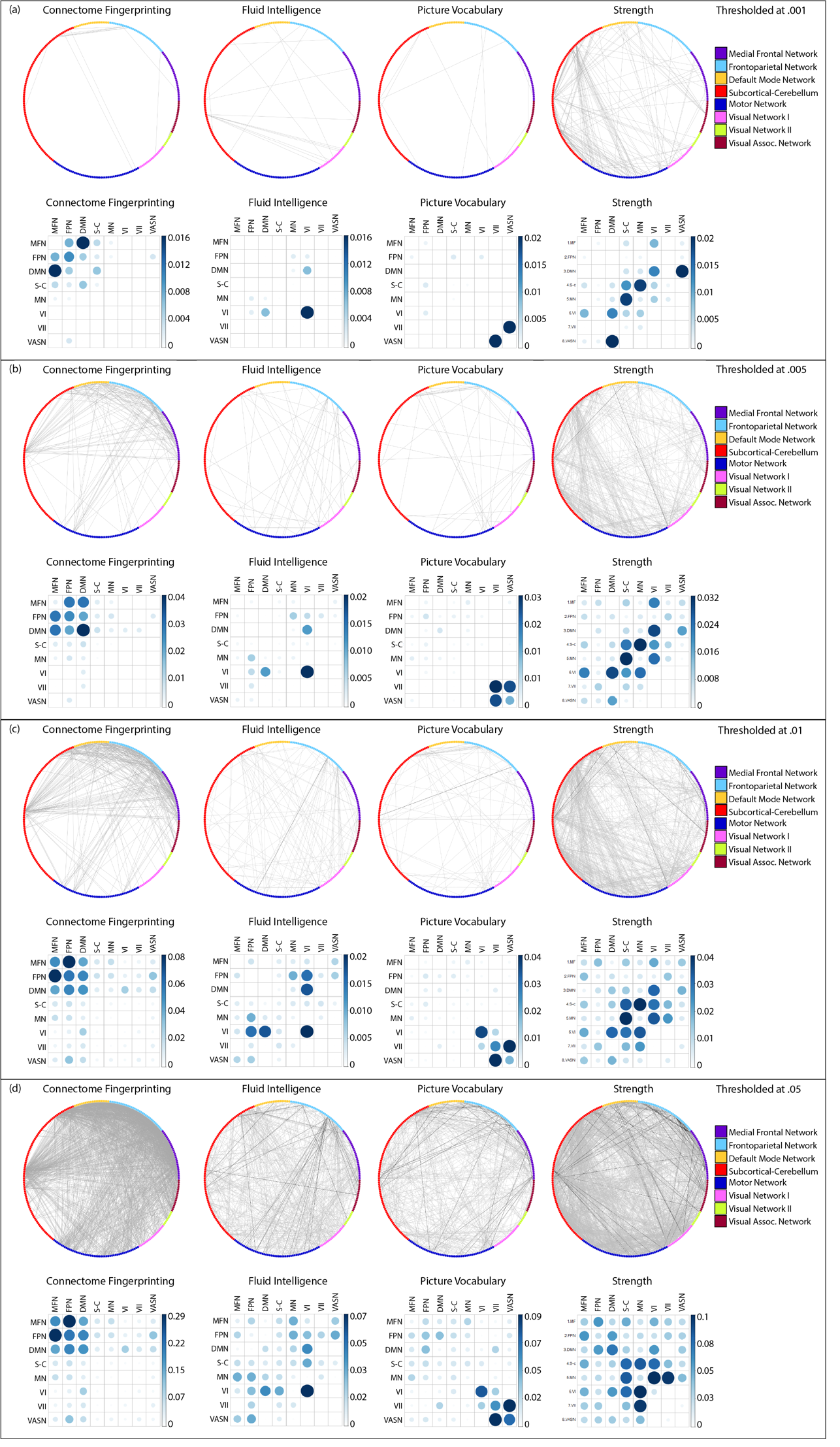

**Supplementary Figure 3:** Single-edge and between-network overlap for fingerprints and the negative prediction models, thresholded at the 99.9^th^, 99.5^th^, 99^th^ and 95^th^ percentile.
